## Supplemental Figures for "Distinct Synaptic Transfer Functions in Same-Type Photoreceptors"

### SUPPLEMENTARY MATERIALS

*Schroeder et al. 2021*

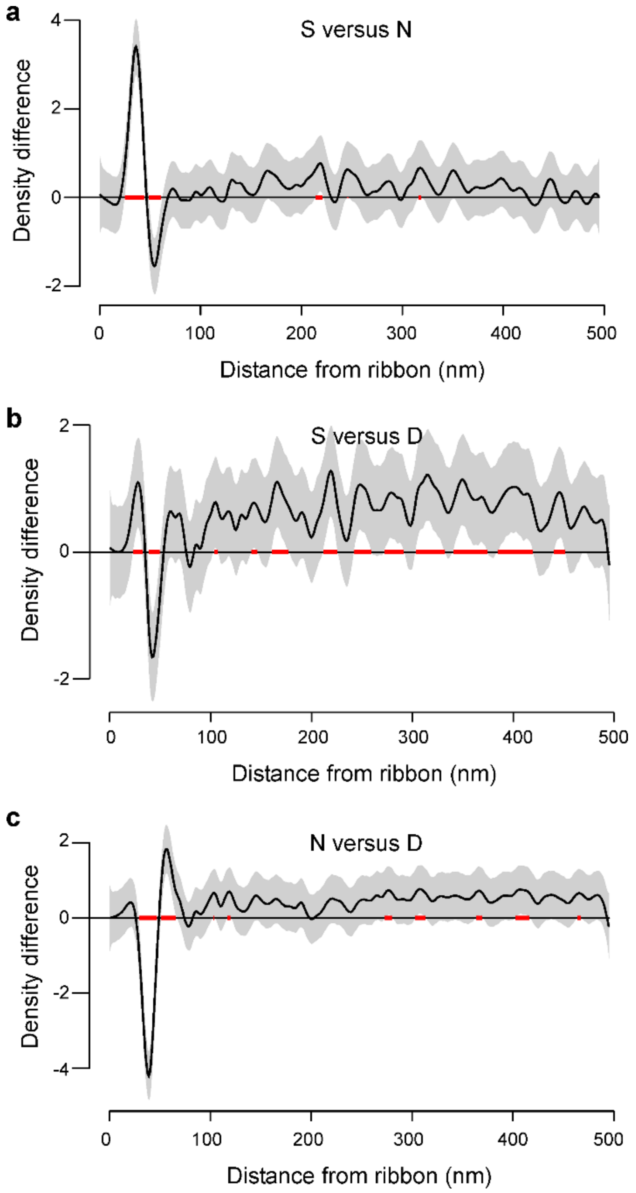


***Figure S1 Statistical comparison of spatial vesicle distributions. a-c,*** *Pair-wise comparison of the three regions, modelled as General Additive Model (Methods). The estimated mean difference (black) and 95% confidence intervals (grey) are shown and distances with significant differences (α=0.05) are highlighted (red).*

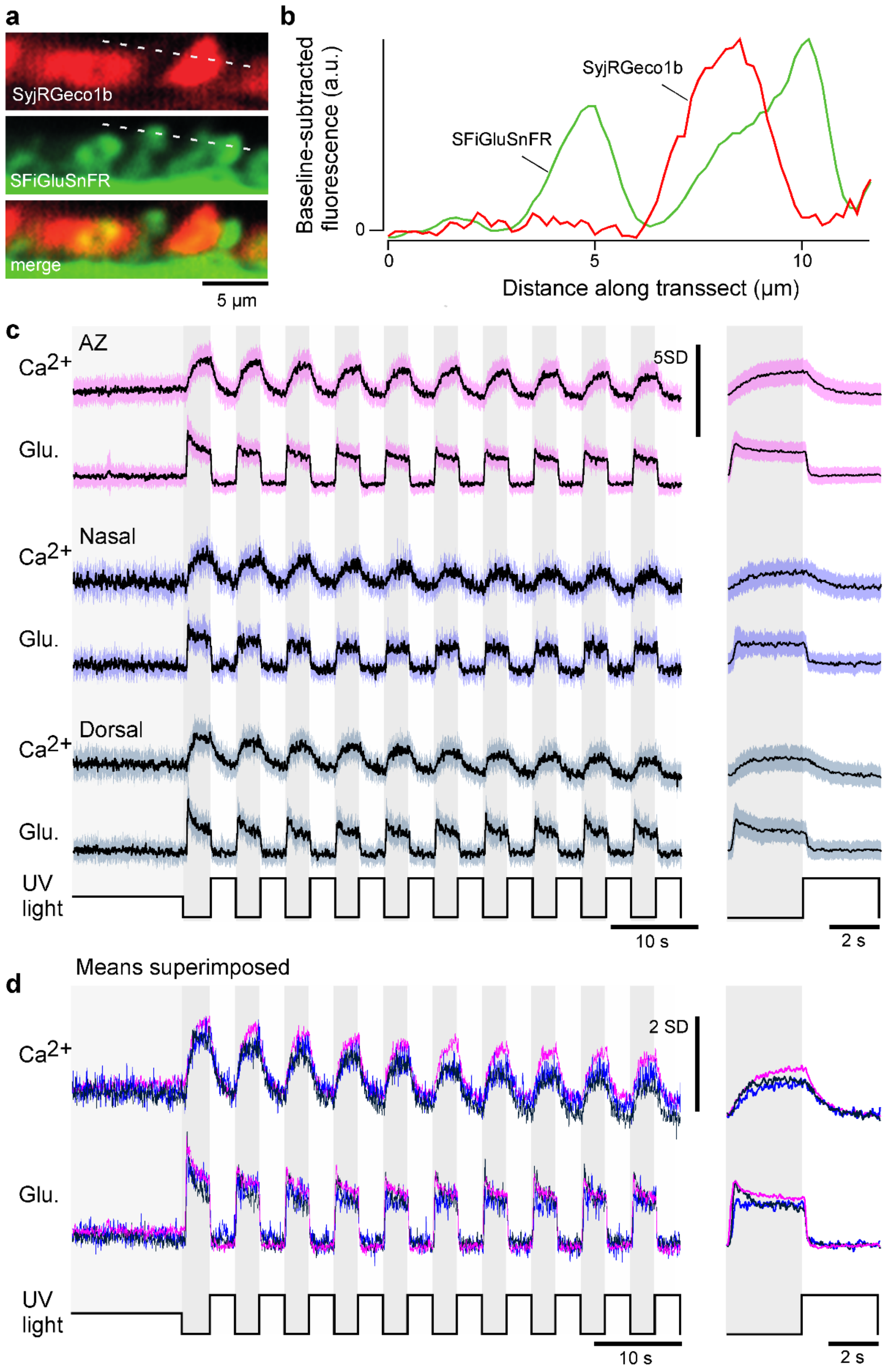


***Figure S2. Mean calcium and glutamate traces per region.*** ***a****, as Fig. 2a, example "red" (top) and "green" (middle) fluorescence channels acquired simultaneously, and merged (bottom) and* ***b****, extracted fluorescence profiles in the two channels as indicated. Note the absence of any obvious "bleed-through", indicating good spectral separation of the two fluorescence detection bands.* ***c,*** *Z-scored recordings before scaling and de-noising (mean±SD).* ***d,*** *Superimposed means of the data shown in (c).*

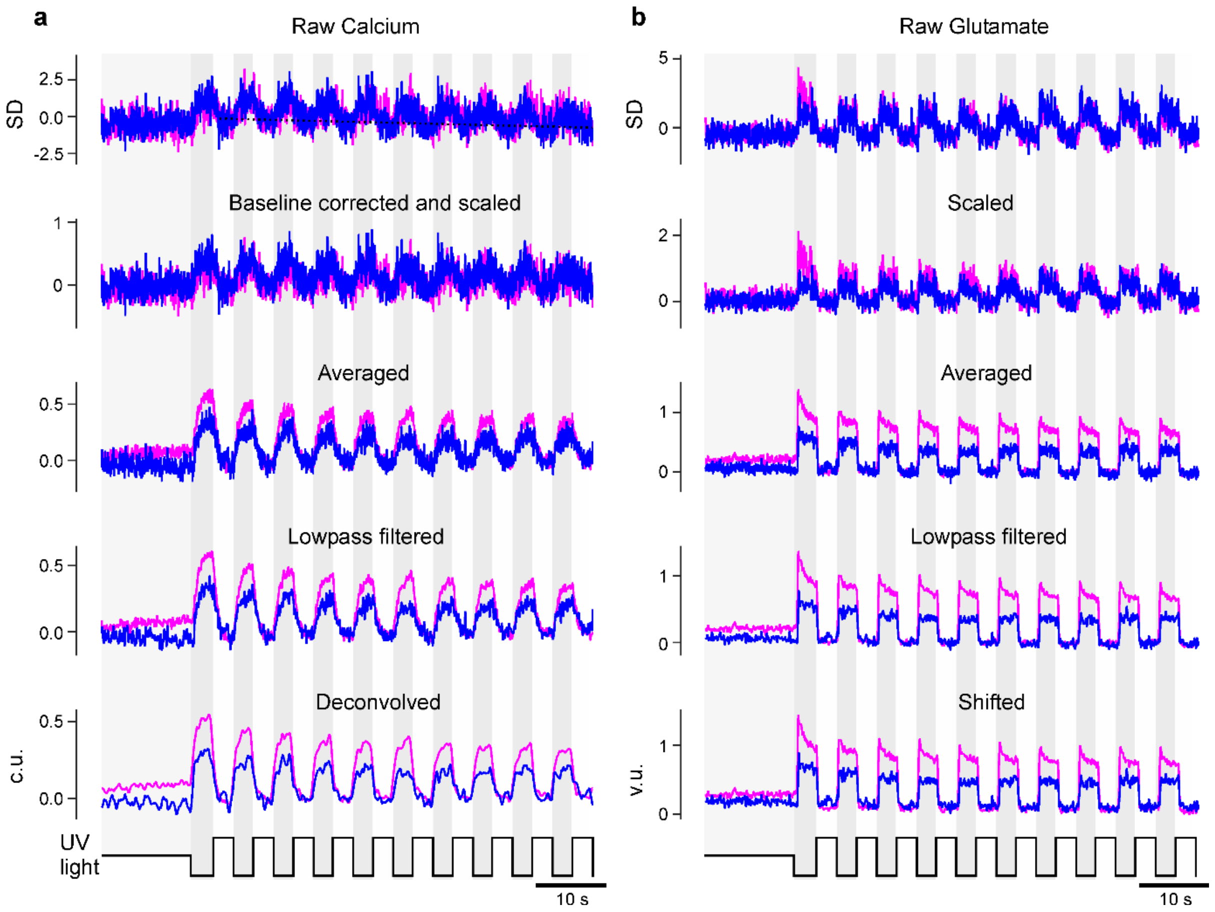


***Figure S3. Pre-processing of calcium and glutamate recordings. a,*** *The raw (z-scored) calcium data is first corrected for a linear decay of the baseline (dotted black line). It is then scaled such that the UV-bright intervals have mean zero and standard deviation one (second row). After averaging all recordings of one zone (third row) the traces are then lowpass filtered and finally deconvolved (last row). We only included the AZ and nasal zone in this illustration.* ***b,*** *As (a) but for the glutamate data, but no baseline correction and deconvolution was applied. As a final step the glutamate traces were shifted to have only non-negative values (last row).*

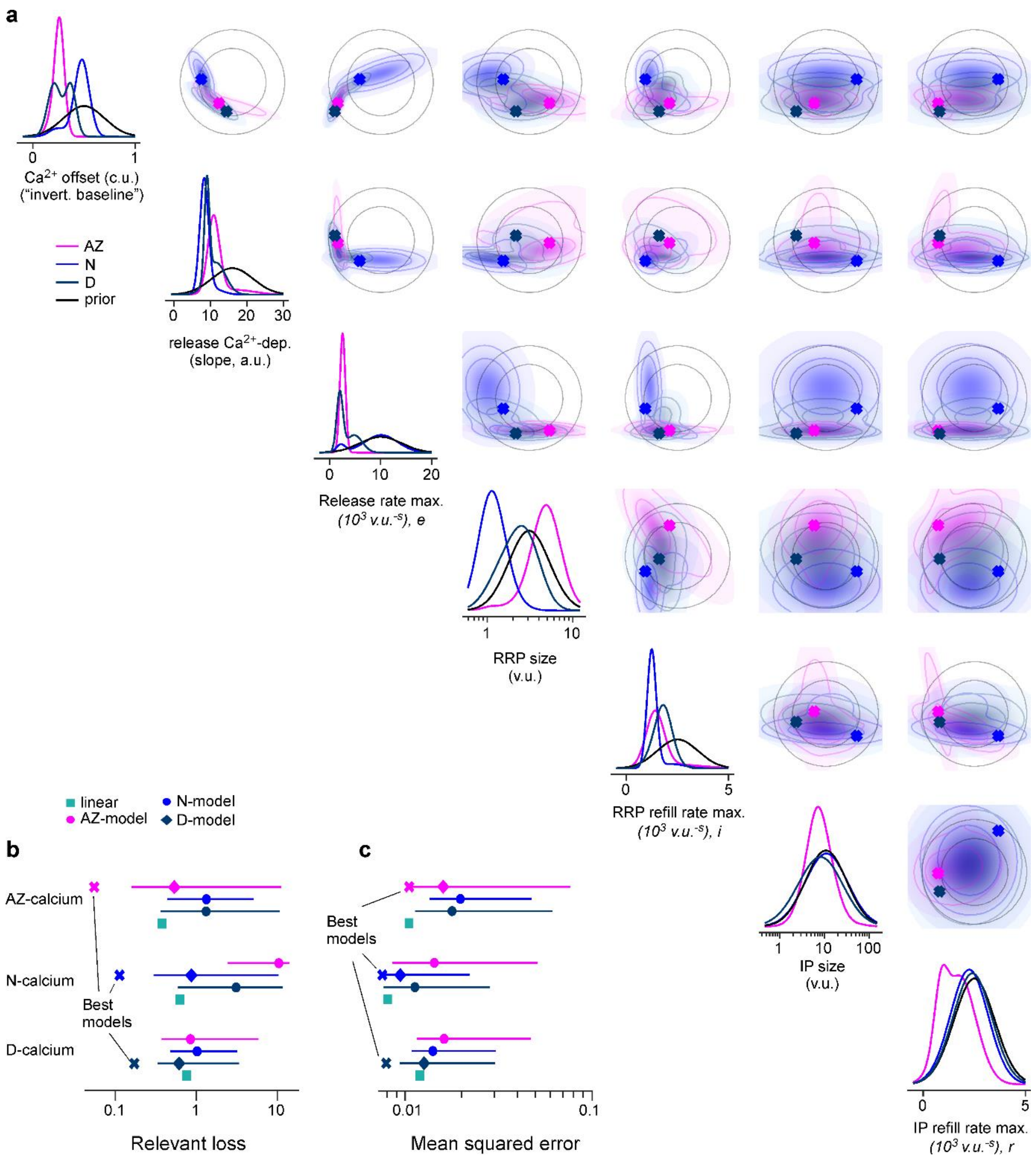


***Figure S4. a,*** *One- and two-dimensional marginals of the posterior as well as of the prior. Best parameters per region are marked as a cross.* ***b,*** *Median with 5$^{th}$ and 95$^{th}$-percentiles (error bars) for the relevant loss based on the summary statistics (Methods), as well as the loss for the best zone specific and the linear baseline model for comparison.* ***c,*** *Same as (b) but loss as Mean Squared Error (MSE) between glutamate recordings and model output.*
